# Immediate and long-term effects of transcranial direct-current stimulation in the mouse primary somatosensory cortex

**DOI:** 10.1101/2020.07.02.184788

**Authors:** Carlos A. Sánchez-León, Isabel Cordones, Claudia Ammann, José M. Ausín, María A. Gómez-Climent, Alejandro Carretero-Guillén, Guillermo Sánchez-Garrido Campos, Agnès Gruart, José M. Delgado-García, Guy Cheron, Javier F. Medina, Javier Márquez-Ruiz

## Abstract

Transcranial direct-current stimulation (tDCS) is a non-invasive brain stimulation technique consisting in the application of weak electric currents on the scalp. Although previous studies have demonstrated the clinical value of tDCS for modulating sensory, motor, and cognitive functions, there are still huge gaps in the knowledge of the underlying physiological mechanisms. To define the immediate impact as well as the after-effects of tDCS on sensory processing, we first performed electrophysiological recordings in primary somatosensory cortex (S1) of alert mice during and after administration of S1-tDCS, and followed up with immunohistochemical analysis of the stimulated brain regions. During the application of cathodal and anodal transcranial currents we observed polarity-specific bidirectional changes in the N1 component of the sensory-evoked potentials (SEPs) and associated gamma oscillations. Regarding the long-term effects observed after 20 min of tDCS, cathodal stimulation produced significant after-effects including a decreased SEP amplitude for up to 30 min, a power reduction in the 20-80 Hz range and a decrease in gamma event related synchronization (ERS). In contrast, no significant long-term changes in SEP amplitude or power analysis were observed after anodal stimulation except for a significant increase in gamma ERS after tDCS cessation. The polarity-specific differences of these long-term effects were corroborated by immunohistochemical analysis, which revealed an unbalance of GAD 65-67 immunoreactivity between the stimulated vs. non-stimulated S1 region only after cathodal tDCS. These results highlight the differences between immediate and long-term effects of tDCS, as well as the asymmetric long-term changes induced by anodal and cathodal stimulation.

**Significance Statement:** Here we provide a first glimpse at the immediate and long-term impact of tDCS on neural processing in alert animals. The obtained results highlight the complexity of tDCS-associated effects, which include both bidirectional as well as asymmetrical modulation depending on the polarity of the stimulation. This asymmetry suggests the implication of different mechanisms underlying the long-term effects induced by anodal and cathodal transcranial currents. Identifying and defining these effects and its associated mechanisms is crucial to help design effective protocols for clinical applications.

## Introduction

Transcranial direct-current stimulation (tDCS) is a safe and well tolerated neuromodulatory technique (Jackson et al., 2017) that relies on the application of constant weak electrical currents on the scalp during several minutes through strategically positioned electrodes (Nitsche and Paulus, 2000; Woods et al., 2016). Given its ability to modulate neuronal excitability, tDCS has attracted the attention of basic and clinical neuroscientists that have investigated its potential to modulate brain function (Nitsche et al., 2008) and treat a variety of neurological conditions such as epilepsy (Regner et al., 2018), attention deficit hyperactivity disorder (ADHD) (Salehinejad et al., 2019) or ataxia (Grimaldi et al., 2014) among others (for a review see Brunoni et al., 2012; Stagg et al., 2018; Miterko et al., 2019).

From a mechanistic point of view, the effects of tDCS on the cortical excitability can be separated into immediate and long-term changes. Immediate effects, appearing at the very moment of electric field application, are related to changes in membrane polarization caused by redistribution of charges in the cells in presence of the externally applied electric field (Chan et al., 1988; Bikson et al., 2019b). On the other hand, long-term effects observed after current cessation require several minutes of stimulation to develop and involve plasticity mechanisms (Huang et al., 2017). Recently, *in vitro* models have been successfully used to show that different neuronal features such as the orientation of the somatodendritic axis with respect to the electric field (Bikson et al., 2004), the neuronal morphology (Radman et al., 2009) or the axonal orientation (Kabakov et al., 2012) are crucial to determine the overall immediate neuronal modulation, showing that purely depolarizing or purely hyperpolarizing stimulation does not exist (Liu et al., 2018). In addition, animal and human studies have revealed that GABA levels (Stagg et al., 2009; Bachtiar et al., 2018; Patel et al., 2019), glial cells (Monai et al., 2016), neurotrophic BDNF (Ranieri et al., 2012) and different receptors such as NMDA (Fritsch et al., 2010), mGluR5 (Sun et al., 2016), AMPA (Stafford et al., 2018; Martins et al., 2019) and adenosine (Márquez-Ruiz et al., 2012) are involved in the long-term effects observed after tDCS. Thus, despite the simplicity of the technique, understanding the overall effect of transcranial electrical currents on brain tissue requires a comprehensive integration of several factors.

Previous tDCS studies have shown its ability to modulate the amplitude and synchronicity of different EEG and LFPs frequencies in human subjects (Antal et al., 2004; Reinhart et al., 2015; Wiesman et al., 2018) and more specifically, to modulate the amplitude of sensory-evoked potentials (SEPs) in both humans (Matsunaga et al., 2004; Dieckhöfer et al., 2006) and animals (Márquez-Ruiz et al., 2012). SEPs are event-related potentials (ERPs) evoked by sensory stimulation (Woodman, 2010) and can be recorded in the human (Sugawara et al., 2015; Vaseghi et al., 2015) and rodent (Castro-Alamancos and Bezdudnaya, 2015) primary somatosensory cortex (S1), constituting a useful test bed for translational studies (Modi and Sahin, 2017; Sánchez-León et al., 2018). The current study looks at both immediate and long-term polarity specific effects, addressing the precise electrophysiological and molecular changes caused by tDCS in the behaving mouse brain.

In this work, we measured the immediate and long-term impact of tDCS applied to the mouse primary somatosensory cortex (S1-tDCS). To assess whether the effects of the stimulation were polarity-dependent (Nitsche and Paulus, 2000), we delivered either anodal or cathodal current. The neuromodulatory effects of S1-tDCS at the electrophysiological level were examined by recording the spontaneous LFPs and sensory-evoked potentials elicited by whisker electrical stimulation in S1 of alert mice. To identify molecular changes induced by S1-tDCS, we performed a poststimulation immunohistochemical analysis of GAD 65-67 and vGLUT1 immunoreactivity in the stimulated brain region.

## Methods

### Animals

Experiments were carried out on adult males C57 mice (University of Seville, Spain) weighing 28-35 g. Before and after surgery, the animals were kept in the same room but placed in independent cages. The animals were maintained on a 12-h light/12-h dark cycle with continuously controlled humidity (55 ± 5%) and temperature (21 ± 1 °C). All experimental procedures were carried out in accordance with European Union guidelines (2010/63/CE) and following Spanish regulations (RD 53/2013) for the use of laboratory animals in chronic experiments. In addition, these experiments were submitted to and approved by the local Ethics Committee of the Pablo de Olavide University (Seville, Spain).

### Surgery

Animals were anesthetized with a ketamine–xylazine mixture (Ketaset, 100 mg/ml, Zoetis, NJ., USA; Rompun, 20 mg/ml, Bayer, Leverkusen, Germany) at an initial dosage of 0.1 ml/20 g. Under aseptic conditions, an anteroposterior (AP) incision in the skin along the midline of the head, from the front leading edge to the lambdoid suture, was performed. Subsequently, the periosteum of the exposed surface of the skull was removed and washed with saline. The animal’s head was correctly positioned to mark the position of bregma as stereotaxic zero. For tDCS administration, a custom-made silver ring electrode, which acted as the active electrode for tDCS, was placed over the skull centered on the right S1 vibrissa area (AP = – 0.9 mm; Lateral = −3 mm; relative to bregma (Paxinos and Franklin, 2004)) (Fig. 1A). To fabricate the silver ring electrodes a silver wire (ø: 635 μm; A-M Systems, WA., USA) was cut into pieces of 1 cm length and one end was curved and welded to form a close loop. Subsequently, the ring was pressed with pliers to flatten the surface creating a 2.5 mm inner ø, 3.5 mm outer ø stimulation surface and was chlorinated. The resulting electrode was introduced into a flexible silicone tubing (inner ø: 0.508 mm; outer ø: 0.939 mm; A-M Systems) for isolating and soldered to a connector pin. Once placed in the correct coordinates, the ring electrode was covered with dental cement (DuraLay, Ill., USA) without pouring it between the electrode and the skull. After that, a hole (2 mm ø) was drilled in the parietal bone inside the ring electrode to expose S1 and the dura mater surface was protected with wax bone (Ethicon, Johnson & Johnson, NJ., USA). In addition, a silver electrode was also implanted over the dura surface under the left parietal bone (AP = - 0.9 mm; Lateral = + 3 mm; relative to bregma (Paxinos and Franklin, 2004)) as electrical reference for the electrophysiological recordings. Regarding the histological experiments where the brains had to be processed for posterior immunohistochemical analysis, the active electrode for tDCS was a polyethylene tubing (inner ø: 2.159 mm; outer ø: 0.325 mm; A-M Systems) placed over the stimulated region. No trepanation was made in the histological experiments to avoid tissue damage. Finally, a head-holding system was implanted, consisting of three bolts screwed to the skull and a bolt placed over the skull upside down and perpendicular to the frontal plane to allow for head fixation during the experiments. The complete holding system was cemented to the skull.

**Figure 1.**
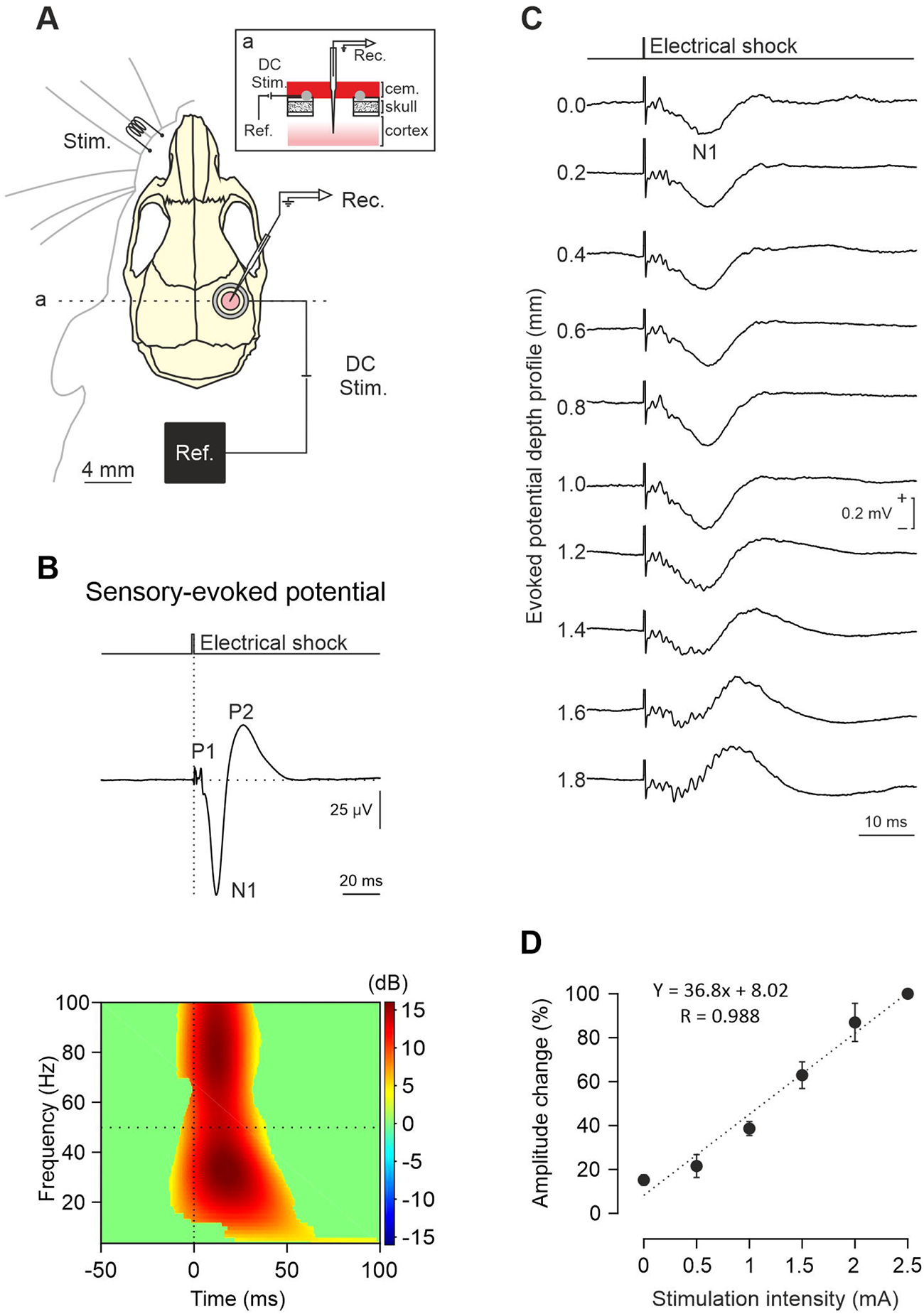
SEP characterization. *A*) Experimental preparation for concurrent tDCS and *in vivo* electrophysiological recordings in S1. *B*) (Up) Waveform showing the different components (P1, N1 and P2) and (down) ERSP of the SEP induced by whisker electrical stimulation (n = 10 mice). *C*) S1-SEP depth profile. Every trace corresponds to an average of 10 SEPs recorded at different depths for a representative mouse. *D*) Quantification of the amplitude change in N1 component of SEPs regarding intensities applied to whisker electrical stimulation. Data normalized with respect to maximum amplitude recorded at 2.5 mA (n = 3 mice).

### Recording and stimulation procedures

Recording sessions began at least two days after surgery. The animals were placed over a treadmill with an infrared sensor for locomotion activity monitoring and the head was fixed to the recording table by means of the implanted head-holding system. Bone wax was removed with the aid of a surgical microscope (SMZ-140, Motic, Barcelona, Spain) and the cortical surface was carefully cleaned.

To stimulate the whiskers, an electrical shock (0.2 ms square pulse, > 2.5 mA) was subcutaneously delivered by a pair of flexible steel electrodes (strand ø: 50.8 μm; coated ø: 228.6 μm; A-M Systems) inserted with the aid of a small needle (25G) under the skin of the left whisker pad with a separation of 2-3 mm. The electrical stimulus consisted of a single square pulse (0.2 ms; 0.5-2.5 mA) delivered by an isolation unit (CS20, Cibertec, Madrid, Spain) connected to a stimulator device (CS420, Cibertec). To characterize the SEPs, a glass micropipette with a tip diameter between 8-10 μm was filled with 3M NaCl, mounted on a micromanipulator (MO-10, Narishige, Tokyo, Japan) and placed over S1 area. In order to map the SEP, the electrical stimulus was delivered at the whisker pad every 10 ± 2 s, the micropipette was lowered and the current intensity adjusted until the maximum amplitude SEP was achieved. Then, the current intensity of whisker electrical pulses was lowered to elicit a SEP with half of the maximum amplitude to allow the observation of an increase or decrease of its components during and after tDCS intervention. All recordings were obtained with an amplifier (BVC-700A, Dagan corporation, MN., USA) connected to a dual extracellular-intracellular headstage (8024, Dagan corporation).

### tDCS

The different protocols for transcranial currents were designed in Spike2 (Cambridge Electronic Design (CED), Cambridge, U.K.) and sent to a battery-driven linear stimulus isolator (WPI A395, Fl., USA) through an analog output from the acquisition board (CED micro1401-3). tDCS was applied between the ring electrode over S1 and a reference electrode consisting of a rubber rectangle (6 cm^2^) attached to the back of the mouse and moistened with electrogel (Electro-Cap International, OH., USA). To measure the actual voltage changes elicited intracranially, transcranial alternating-current stimulation (tACS) was applied at ± 2, ± 20 and ± 200 μA (± 0.0426, ± 0.426 and ± 4.26 mA/cm^2^) at 1 Hz. Every sinusoid wave lasted for 10 seconds and each intensity was randomly distributed and repeated three times. Glass micropipettes were used to sequentially record at different depths, ranging from the cortical surface until 4 mm below, in 1 mm steps. To characterize the immediate effects induced by tDCS, 15 s pulses of anodal and cathodal tDCS (including 5 s ramp-up and 5 s rampdown) of varying intensities (50, 100, 150 and 200 μA), were applied separated by 10 s of non-stimulation. To index long-lasting changes, tDCS was delivered during 20 min at 200 μA for cathodal stimulation, and for 20 min at 150 μA for anodal stimulation. Differences in the intensity used for cathodal and anodal were due to amplifier noise issues with higher anodal currents. SEPs in response to contralateral whisker stimulation (every 10 ± 2s) were recorded before (control), during (immediate effects), and after (long-term effects) tDCS presentation. For immunohistochemical experiments, the tubing used as tDCS active electrode was filled with electrogel and a metallic electrode from the stimulus isolator was immersed on it.

### Histology

To characterize potential histological changes associated to tDCS, a different set of animals received 20 min of anodal, cathodal or sham tDCS at 200 μA. 15 min after tDCS mice were deeply anesthetized with ketamine–xylazine mixture (Ketaset, 100 mg/ml; Rompun, 20 mg/ml) and perfused transcardially with 0.9% saline followed by 4% paraformaldehyde (PanReac, Barcelona, Spain) in PBS. The brains were removed and stored in 4% paraformaldehyde for 24 hours, cryoprotected in 30% sucrose in PBS the next 48 hours, and then cut into 50 μm coronal slices with a freezing microtome (CM1520, Leica, Wetzlar, Germany). Sections were processed “free-floating” and passed through all procedures simultaneously to minimize differences in immunohistochemical staining. After three washes of 10 min with PBS, sections were blocked with 10% Normal Donkey Serum (NDS, 566460, Merck, Darmstadt, Germany) in PBS with 0.2% Triton X-100 (Sigma-Aldrich, Mo., USA) (PBS-Tx-10% NDS) and then incubated overnight at room temperature in darkness with the primary antibody solution containing mouse anti-vesicular Glutamate Transporter 1 (vGLUT1, 1:1000, MAB5502, Merck) or rabbit anti-Glutamate Decarboxylase 65-67 (GAD 65-67, 1:1000, AB1511, Merck). After three washes, sections were incubated for 1 hour at room temperature in darkness with appropriate secondary antibodies: Alexa Fluor 488 donkey anti-mouse IgG (H+L) (1:400, A21202, Thermo Fisher Scientific, Mass., USA), Alexa Fluor 555 donkey anti-rabbit IgG (H+L) (1:400, A31572, Thermo Fisher Scientific) in PBS-Tx-5% NDS. After three washes with PBS, sections were mounted on glass slides and coverslipped using Dako Fluorescence Mounting Medium (Dako North America, CA., USA). For confocal imaging, an *in vivo* confocal microscope (A1R HD25, Nikon, Tokyo, Japan) was used. Z-series of optical sections (0.5 μm apart) were obtained using the sequential scanning mode.

### Data collection

Recording signal from the amplifier, tDCS converted signals, infrared sensor signals from wheel motion and 1-V rectangular pulses corresponding to whisker electrical stimulation presented during the different experiments were digitally stored on a computer for quantitative off-line analysis (CED micro1401-3). Collected data were sampled at 25 kHz for SEP and LFP recordings, with an amplitude resolution of 16 bits. The remaining non-neuronal signals were sampled at 5 kHz.

#### Data analysis

##### Intracranial electric field analysis

To estimate the electric field strength during tACS, “DC remove” process from Spike2 was applied (time constant of 0.5 s) to correct for possible baseline drifts unrelated to stimulation and set the channel offset to zero. From every sinusoid wave, the peak-to-peak value (electric potential) from the LFP evoked by transcranial electrical stimulation (tES) was measured and averaged for a given intensity and depth. Finally, for each intensity the electric field strength (differences between potentials) was calculated by computing the difference in peak-to-peak values between two consecutive depths (1 mm in distance).

##### Peak-to-peak amplitude and latency

SEP amplitude was computed by the peak-to-peak command in Spike2 software, where the maximum negative voltage value (N1) was subtracted from the maximum positive voltage value (P1) of the preceding peak. SEP latency was determined as the time from whisker stimuli to the maximum N1 peak value. SEPs recorded when the animal was running were removed from the analysis (given the dramatic decrease in some SEP’s components due to the movement), as well as those potentials presenting electrical artifacts.

##### tDCS immediate effects analysis

For tDCS immediate effect experiments, SEPs induced by left whisker-pad stimulation were recorded 1 s before tDCS ramp-up (control) and 1 s before tDCS ramp-down (immediate effects). For every tDCS trial, amplitude values during tDCS were normalized (dividing them by the immediately preceding control value multiplied by 100), and latency values were normalized subtracting them by the latency value of the preceding control SEP. Data obtained from the same tDCS polarity and intensity were averaged and compared between animals.

##### tDCS long-term effects analysis

For tDCS long-term effect experiments, SEPs induced by left whisker-pad stimulation (delivered every 10 ± 2 s) were recorded 20 min before (control), 20 min during, and 60 min after tDCS. Every 5-minute interval SEP waveforms were averaged and, for comparison between animals, amplitude values were normalized dividing them by the mean of the baseline values (control condition, before tDCS) multiplied by 100. Using this normalization, the baseline values were always close to 100 % but the variance is the same as in the raw data. For latency normalization, latency values before, during and after tDCS were subtracted by the mean of the baseline (control period) values.

##### Fluorescence immunohistochemistry

Confocal images were processed in ImageJ (https://imagej.nih.gov/ij/) with the image-processing package Fiji (http://fiji.sc/Fiji) using a custom built macro. To subtract fluorescence background noise five square regions of interest (ROI) of 30 x 30 pixels (26.22 μm^2^) were placed over unlabeled nuclei in each image, and the obtained maximum brightness average was set as the minimum value for “setThreshold”, so the pixels with values lower than the average were considered as non-fluorescent. Then, a copy of the image was converted to binary to visually validate the procedure, and the complete process was repeated until the threshold properly discriminated our signal from the noise. To analyze particles, five square ROI of 100 x 100 pixels (291.31 μm^2^) were randomly placed over regions absent of nuclei or unspecific noise (as for example blood vessels). Each image inside the ROI was converted to binary and the “Analyze Particles” command was used to count and measure aggregates of vGLUT1 and GAD 65-67. Particles were sorted as small (size = 10-25), medium (size = 26-46) or big (size = 47-100) and averaged to obtain one value per hemisphere per animal.

##### Event-related potential (ERP) analysis

ERP analysis was performed in EEGLAB rev.14.1.2 toolbox using Matlab 2015a software package. Data were segmented in intervals of 1000 ms, from 500 before to 500 ms after the electrical stimulation of the whisker. The baseline was corrected by subtracting the mean voltage level in the first 500 ms interval of the window. To eliminate the artifacts in the original signal produced by electrical stimulation in the whiskers, the algorithm developed by Feuerstein et al. (2009) was used, consisting in the identification and characterization of the types of noise, and their subsequent subtraction from the otherwise unprocessed data set. For each condition (cathodal, anodal and sham), temporal period (20 minutes control, 20 minutes during tDCS, 20 minutes post-tDCS and 40 minutes post-tDCS) and subject, data were averaged to obtain the SEP by using the electrical stimulation as a trigger. The obtained SEPs during the different temporal periods were statistically compared using parametric tests in EEGLAB toolbox.

To analyze the spectral dynamics of the neural oscillations (Fast Fourier Transform - FFT) and Event-Related spectral perturbation (ERSP), an analysis of the *induced activity* was performed. For the isolation of induced from evoked activity, the average SEP from every subject was subtracted from each condition (cathodal, anodal and sham), temporal period (20 minutes control, 20 minutes during tDCS, 20 minutes post-tDCS and 40 minutes post-tDCS) and subject (Tallon-Baudry and Bertrand, 1999; Bastiaansen and Hagoort, 2003).

##### Fast Fourier transform (FFT)

FFT analyses was performed in EEGLAB rev.14.1.2 toolbox using Matlab 2015a software package. FFT from every subject was extracted from each condition (cathodal, anodal and sham) and temporal period (20 minutes control, 20 minutes during tDCS, 20 minutes post-tDCS and 40 minutes post-tDCS), as well as the average FFT for all subjects in each condition and temporal period. Subsequently, FFTs from 20 minutes post-tDCS and 40 minutes post-tDCS were compared with the FFT from 20 minutes of control condition for every condition independently. A statistical analysis by permutations (p < 0.05) with a false discovery rate (FDR) for multiple comparisons was applied.

##### Event-related spectral perturbation (ERSP) analysis

A time-frequency signal analysis was performed trial-by-trial using Hanning-windowed sinusoidal wavelets at 1 cycle (lowest) to 13.3 cycles (highest). Changes in event-related dynamics of the signal spectral power were studied using the ERSP index (Makeig, 1993). ERSP quantifies the mean change in spectral power (dB) from the baseline at different latencies and frequencies with respect to the event (electrical stimulation of the whisker). Significance thresholds for ERSP were calculated by a bootstrap distribution (p < 0.05), extracted randomly from the baseline data (from −330 ms to 0 ms) and applied 400 times (Makeig et al., 2002). Additionally, the ERSP of the different temporal periods were statistically compared by permutations analysis (p < 0.05). To visualize changes between temporal periods, differential activity (DA) was computed by subtracting the ERSP of the control period to the ERSP of each period. Thus, warm colors in DA mean that spectral power was higher in that period with respect to control period, and cooler colors mean lower spectral power with respect to control period.

### Statistical analysis

SigmaPlot 11.0 (Systat Software Inc, San Jose, CA., USA), IBM SPSS version 25 (IBM, Armonk, NY)) and Matlab 2015a (MathWorks Inc.) were used for statistical analysis. For immediate effects experiments, statistical significance of differences between groups was inferred by a two-way repeated-measures analysis of variance (ANOVA), with CURRENT INTENSITY (50, 100, 150 or 200 μA) and POLARITY (anodal or cathodal) as within-subject factors, and the post hoc Holm-Sidak test for multiple comparisons. For long-term effects experiments, a two-way repeated-measures ANOVA was performed to infer statistical differences with TIME (temporal periods of 5 minutes each: one time point for control, four time points during tDCS/sham and twelve time points after tDCS/sham) as within-subject factor, and tDCS POLARITY (anodal, cathodal or sham) as between-subjects factor. The post hoc Bonferroni test was applied for multiple comparisons. For immunohistochemical experiments, statistical comparison for fluorescence levels was inferred by a two-way mixed ANOVA with BRAIN HEMISPHERE (non-stimulated vs. stimulated hemisphere) as within-subject factor and tDCS POLARITY (anodal, cathodal or sham) as between-subjects factor. The post hoc Bonferroni test was applied for multiple comparisons. The results are shown as mean ± SEM. Statistical significance was set at p < 0.05 in all cases.

## Results

### Characterization of sensory-evoked potentials in response to whisker stimulation

To index potential changes in the neuronal excitability of S1 during and after tDCS, SEPs in response to whisker stimulation were chronically recorded in alert head-restrained mice (n = 10; Fig. 1A). Electrical whisker stimulation evoked a contralateral short-latency SEP in the vibrissa S1 area (Fig. 1B) consisting of a first positive component (P1) peaking at 3.8 ± 0.2 ms (n = 10), followed by a negative wave (N1) at 12.6 ± 1.2 ms (n = 10), and finally a positive slower component (P2) peaking at 26.2 ± 2.8 ms (n = 10). The amplitude and latency of the N1 component of SEP varied along the recording sites across cortical layers (Fig. 1C), reaching maximum amplitude between 0.8 – 1.0 mm depth and showing a polarity inversion at deeper recording sites. During experimental sessions depth profiles were obtained from all the participating mice selecting those recording sites where the amplitude of N1 was maximum. The final amplitude of the N1 component was linearly depended on the intensity of the electrical stimuli applied to the whiskers, as shown in figure 1D (R = 0.988; p < 0.001; n = 3). Finally, the ERSP of SEPs (Fig. 1B, at the bottom) was characterized by a significant increase in power spectrum for all analyzed frequencies (3-100 Hz) associated with the first 50 ms of SEP after the whisker electrical shock. As observed in figure 1B, two major frequency bandwidths were maximally enhanced, one at 20-40 Hz and other at 60-100 Hz.

### tACS-elicited electric field decays with distance from the active electrode

In a first experiment, we determined the actual electric field gradient along the brain tissue imposed by transcranial electrical stimulation (tES) application in our experimental design. Animals (n = 6) were prepared for chronic recording of LFPs in the S1 area in alert condition during simultaneous application of low-frequency tACS (1 Hz) (Fig. 2A). Recordings were performed by inserting a glass micropipette (1-5 MΩ of impedance, lateral angle 20°) through a ring electrode previously implanted on the mouse skull (Fig. 1A, inset). Differential recordings were obtained between the glass micropipette and a silver reference electrode placed over the dura and chronically implanted in the previously performed surgery (Ref. in Fig. 2A). Differential recordings were sequentially performed every 1 mm from the cortical surface to 4 mm depth. Figure 2B shows the grand average obtained from recordings at different depths including the data from all six animals. Electric fields were determined by subtracting the generated electric potentials during 1 Hz tACS applied in randomly interleaved trials at three intensities (2, 20 and 200 μA). The calculated electric fields at different depths and intensities for all animals are represented in figure 2C. Under the active electrode, the magnitude of the electric field decreased with depth in a logarithmic manner for the three tested intensities (data are presented with logarithmic abscissa axis for visual facilitation, Fig. 2C). To calculate the electric field imposed by tES at different intensities (50, 100, 150 and 200 μA) in the recording site (~1 mm) we used a linear regression equation extracted from the relation between tACS intensity and electric field strength (E = −0.4473 * I - 0.731; R = 0.997; p = 0.0028; n = 3). The calculated electric field strength induced by 50, 100, 150 and 200 μA at the recording site was 23.1, 45.5, 67.8 and 90.2 V/m, respectively.

**Figure 2.**
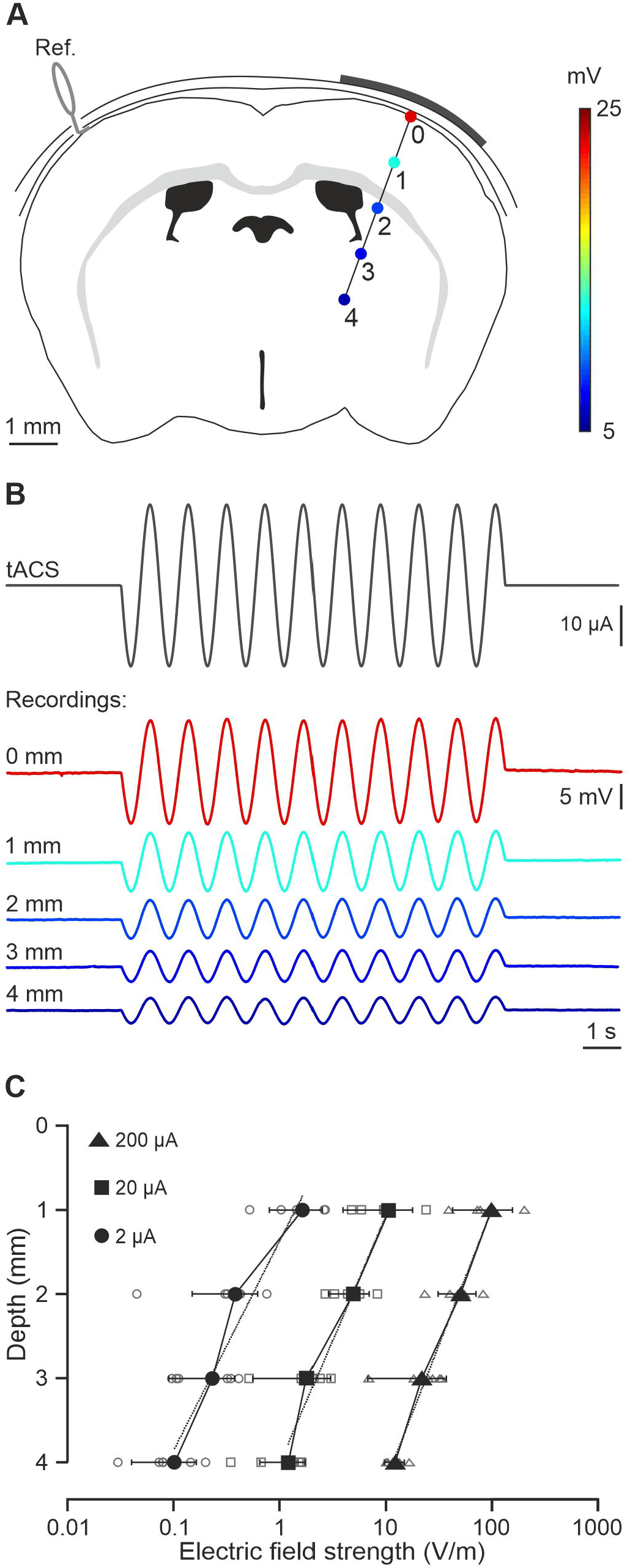
Intracranial electric fields induced by S1-tDCS. *A*) Schematic representation of electric potentials recorded in S1 at different depths. *B*) tACS stimulation (top trace) applied over the scalp and grand average (n = 6 mice, unprocessed data) of the actual potentials generated at different depths (from 0 to 4 mm). *C*) Average (filled symbols) and individual (empty symbols) electric fields recorded at different depths for ± 2 (circles), ± 20 (squares) and ± 200 μA (triangles) tACS.

### tDCS increases and decreases the amplitude of simultaneously recorded SEPs in a polarity and intensity-dependent manner

To test the immediate effects of tDCS on S1 excitability we recorded SEPs induced by whisker pad stimulation during simultaneous short-duration (15 s, including 5 s ramp up and 5 s ramp down (Fig. 3A)) anodal and cathodal tDCS pulses, at 4 randomly interleaved intensities (50, 100, 150 and 200 μA). SEPs recorded just before tDCS pulses were used as controls, to normalize the peak-to-peak amplitude in anodal and cathodal conditions. As a result, the amplitude of N1 component of the SEPs recorded in S1 was significantly increased and decreased in response to anodal and cathodal tDCS, respectively, when compared to the control condition. Figure 3B shows the averaged SEPs (n = 30) during the control condition (black trace), anodal (red trace) and cathodal (blue trace) tDCS applied at 50, 100, 150 and 200 μA conditions for a representative animal. Mean data obtained from the group of animals participating in the experiment (n = 14) are represented in figure 3C. Thus, increasing the intensity of anodal tDCS progressively increased (to a maximum of 141.5 ± 5.3 % at 200 μA) the amplitude of simultaneously recorded SEPs with significant differences for all intensities compared to cathodal situation, whereas increasing the intensity of cathodal tDCS significantly decreased the SEPs amplitude to a minimum of 73.2 ± 4.4 % at 200 μA (two-way repeated-measures ANOVA, CURRENT INTENSITY: F_3,39_ = 3.316, p = 0.030; POLARITY: F_1,13_ = 51.081, p < 0.001, interaction CURRENT INTENSITY X POLARITY: F_3,39_ = 27.818, p < 0.001; Holm-Sidak, p < 0.05; Fig. 3C). In summary, the magnitude and direction of tDCS effects on the N1 amplitude of simultaneously recorded SEPs were dependent on the applied polarity and intensity.

**Figure 3.**
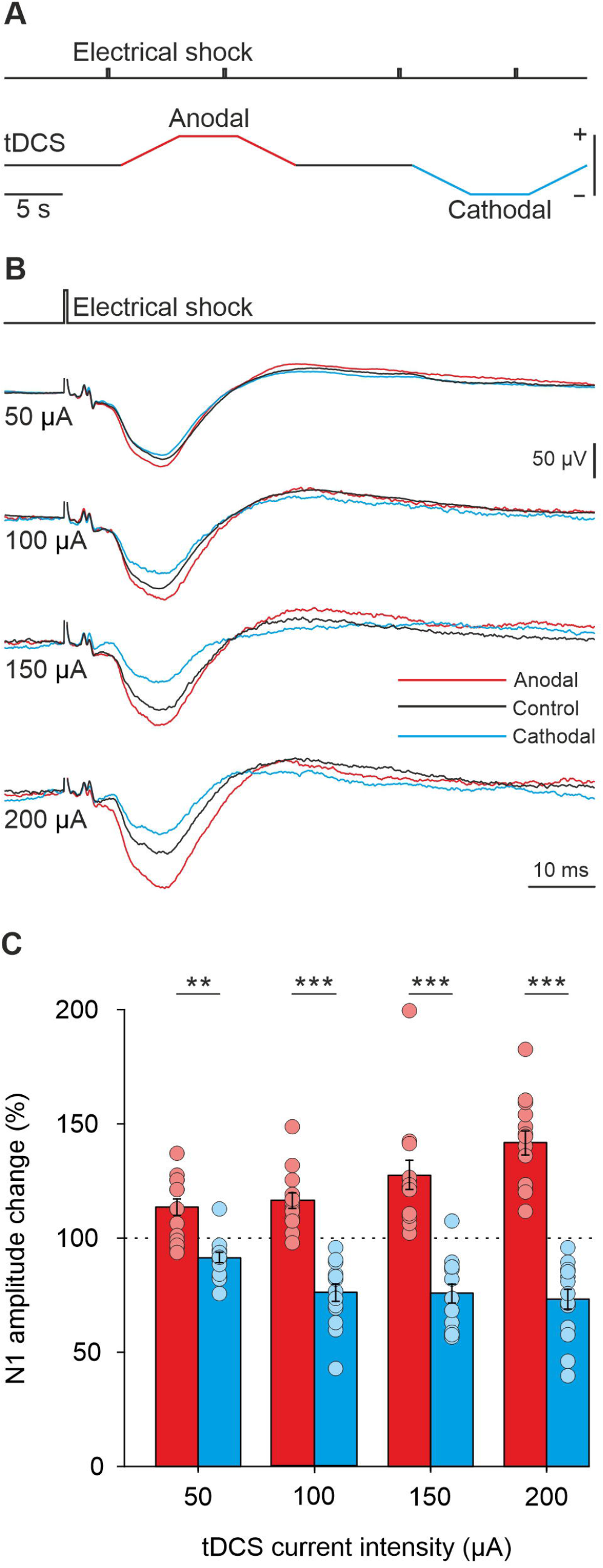
tDCS immediate effects over SEPs in S1 cortex. *A*) Schematic representation of tDCS protocol. *B*) SEP average (n = 30) from a representative animal during control (black trace), anodal (red trace) and cathodal (blue trace) tDCS applied at 50, 100, 150 and 200 μA. *C*) Quantification and statistical results of tDCS effects on SEP amplitude. Mean (bars) and individual amplitude data (circles) are represented as percentage of change with respect to control values for all animals (n = 14 mice). Two-way repeated-measures ANOVA, CURRENT INTENSITY effect, F_3,39_ = 3.316, p = 0.030, POLARITY effect, F_1,13_ = 51.081, p < 0.001, CURRENT INTENSITY X POLARITY interaction, F_3,39_ = 27.818, p < 0.001, Holm-Sidak post hoc test, **p < 0.01; ***p < 0.001.

### tDCS induces asymmetric long-term after-effects on SEP amplitude depending on different current polarity

To test potential long-term effects of tDCS over S1 excitability we recorded SEPs induced by whisker pad stimulation (every 10 ± 2 s) in three different experimental conditions. Animals were prepared for SEP recording and simultaneous tDCS and randomly assigned to the anodal (n = 10), cathodal (n = 10) or sham (n = 10) group. During experimental sessions SEPs were recorded for 20 min before tDCS, during continuous anodal (150 μA, 20 min), cathodal (200 μA, 20 min) or sham (150 μA, 30 s) tDCS, and for 1 hour after tDCS. As observed in figure 4A, tDCS has a significant effect on the normalized N1 amplitude of SEPs for both anodal and cathodal polarity (twoway repeated-measures ANOVA, TIME: F_5.4,146.5_ = 2.792, p = 0.016; POLARITY: F_2.27_ = 20.895, p < 0.001, interaction TIME x POLARITY: F_10.9,140.5_= 8.967, p < 0.001; Bonferroni, p < 0.05). Interestingly, anodal tDCS significantly increased the amplitude of SEPs (up to a maximum of 158.2 ± 11.0 %, n = 10) with respect to control values only during simultaneous tDCS intervention (Bonferroni, p < 0.05; red filled diamonds in Fig. 4A), whereas cathodal tDCS decreased the amplitude of SEPs with respect to control values reaching its maximum effects (maximum of 33.1 ± 7.9 %, n = 10) during simultaneous tDCS application remaining significantly decreased for 25 min after tDCS cessation (Bonferroni, p < 0.05; blue filled squares in Fig. 4A). No significant effects were observed in the amplitude of N1 component of SEPs in the sham group (Bonferroni, p < 0.05; black triangles in Fig. 4A). As expected, significant differences between anodal and sham group were restricted to the tDCS period (Bonferroni, p < 0.05, Fig. 4A, asterisks) whereas significant differences were maintained during and for 35 min after tDCS when comparing the cathodal with the sham group (Bonferroni, p < 0.05, Fig. 4A, asterisks).

**Figure 4.**
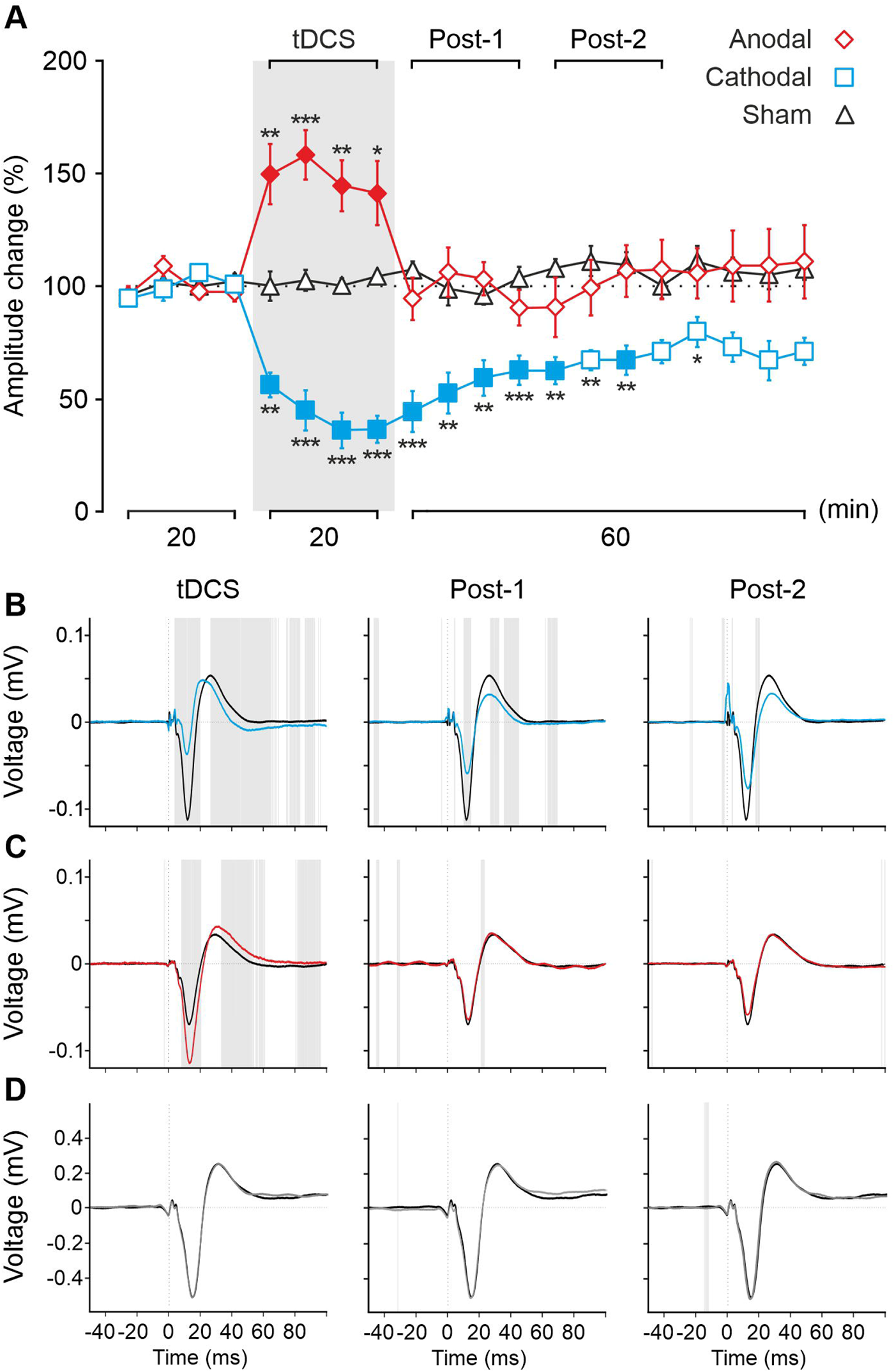
tDCS long-term effects over SEPs in S1 cortex. *A*) Normalized amplitude change of N1 averaged every 5 min for 20 min of anodal (red diamonds), cathodal (blue squares) or sham (black triangles) tDCS. Two-way repeated-measures ANOVA, TIME: F_5.4,146.5_ = 2.792, p = 0.016; tDCS POLARITY: F_2,27_ = 20.895, p < 0.001, interaction TIME x tDCS POLARITY: F_10.9,146.5_= 8.967, p < 0.001. Filled symbols represent statistical differences with the last control period (n = 10 mice, p < 0.05, Bonferroni post hoc test). Asterisks mark statistical differences between the same temporal period for anodal or cathodal with sham tDCS (n = 10 animals, *p < 0.05; **p < 0.01; ***p < 0.001, Bonferroni). *B-D*) ERP analysis comparing all SEPs averaged 20 min before tDCS (black trace) with respect to averaged SEPs during 20 minutes of tDCS (first column), or averaged SEPs in the first 20 min after tDCS cessation (Post-1, second column) or the next 20 min (Post-2, third column) for cathodal (blue traces), anodal (red traces) or sham (grey traces) tDCS. Grey shadow represents statistical differences between points (p < 0.05, paired t-test).

We also analyzed the grand average SEP waveforms induced by whisker pad stimulation (ERP analysis). This analytical approach allowed the exploration of significant differences at every time point of the recorded SEPs. The voltage analysis of the SEP compared the 20 min averaged SEP before, during and after tDCS. As shown in figure 4B (left blue trace), cathodal tDCS significantly decreased the amplitude of different components in the simultaneously recorded SEPs (gray shading indicates p < 0.05, n = 10, Fig. 4B). This effect was progressively reduced after cathodal tDCS cessation, being maintained during the first 20 min (middle Fig. 4B) and almost inexistent for the next 20 min period (right Fig. 4B). On the other hand, anodal tDCS significantly increased the amplitude of different components in the simultaneously recorded SEPs (gray shading indicates p < 0.05, n = 10, Fig. 4C) whereas no remarkable significant effects were observed 20 min (middle Fig. 4C) or 40 min (right Fig. 4C) after anodal tDCS. As expected, no remarkable significant effects were observed in the sham group (n = 10, Fig. 4D).

In addition to the analysis of SEP amplitude we also analyzed the impact of tDCS on the *induced response* where the average SEP from every animal was subtracted from each condition, temporal period and subject. In a first approach, we tested for general changes in the amplitude of neuronal oscillations at different frequencies by FFT analysis of the induced activity. For that, we analyzed the potential significant changes in the power spectrum of the induced activity (selecting a temporal window of ±2.5 s respect of the whisker stimulus) before (PRE) and after (first 20 min: POST1; and next 20 min: POST2) cathodal (Fig. 5A), anodal (Fig. 5B) tDCS and sham condition (Fig. 5C). No CONTROL vs. DURING temporal periods were analyzed in this case because tDCS-associated artifacts were often present in the selected time intervals (5 s duration). Significant differences (n = 10 for anodal and sham, n = 9 for cathodal condition; permutation analysis with FDR for multiple comparisons, p < 0.05) were observed between POST1 and PRE condition in the 20-80 Hz band for cathodal tDCS (Fig. 5A, left column, gray shading indicates p < 0.05) showing a decrease in the amplitude of cortical oscillations in this bandwidth. These differences were not present (except for a few points in the 50-70 Hz range) when PRE and POST2 were compared (Fig. 5A, right column, gray shading indicates p < O.O5). Interestingly, there were no significant changes in any of the comparisons after anodal tDCS (Fig. 5B) or sham condition (Fig. 5C).

**Figure 5.**
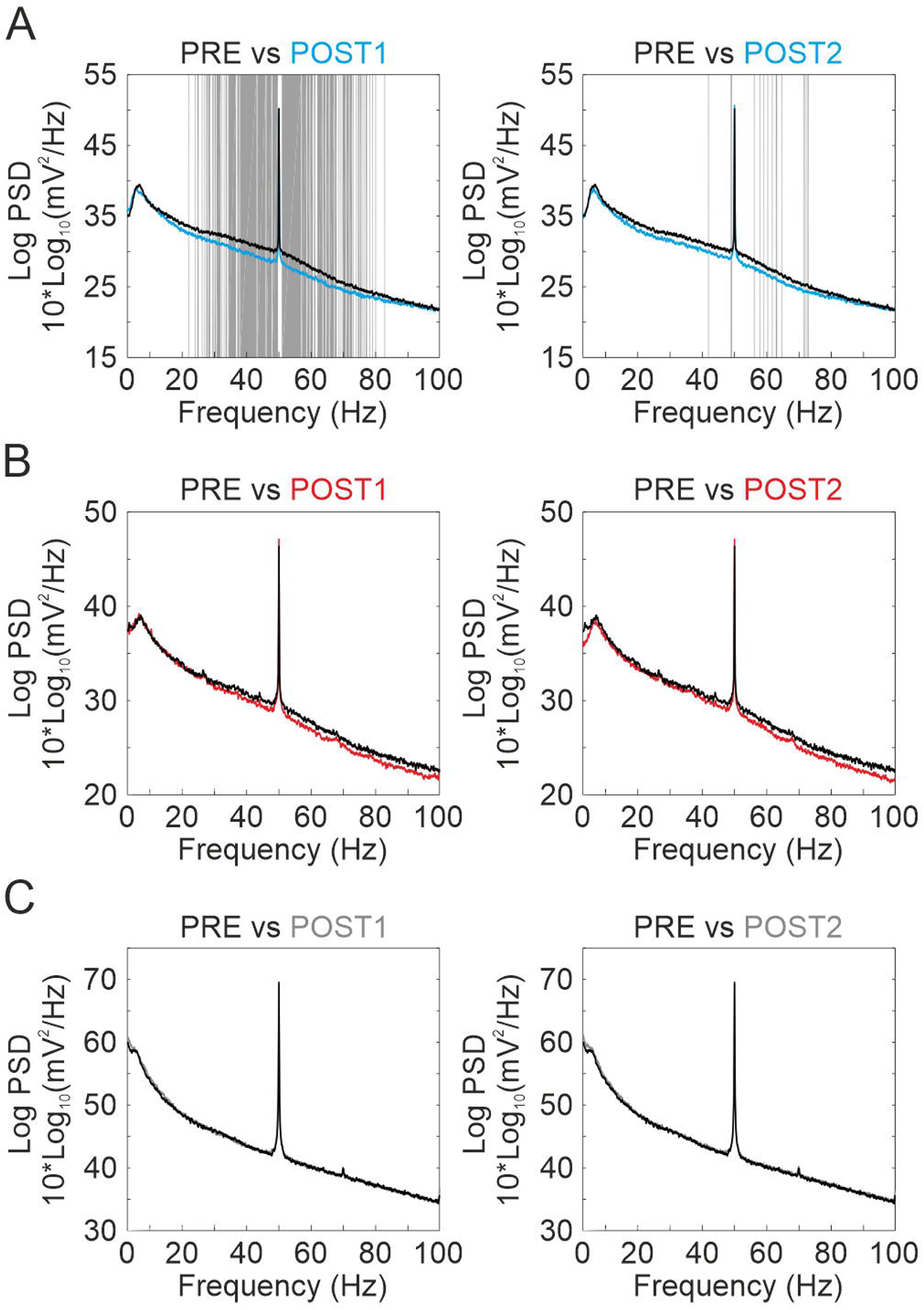
FFT analysis. *A-C*) Comparison between the FFT obtained 20 min before the stimulation and the first 20 min after tDCS cessation (POST1, first column) or the next 20 min (POST2, second column) for cathodal (*A*), anodal (*B*) or sham (*C*) tDCS. Grey shadow represents statistical differences for a specific frequency (p < 0.05, permutations).

To know more about the organization of these frequency differences in the temporal domain we carried out a spectral dynamic analysis of induced response (from 50 ms before to 100 ms after whisker stimulation) associated to sensory stimulation before, during and after tDCS (Fig. 6). We found that during the 20 minutes of cathodal tDCS the spectral power was significantly lower with respect to the control period (permutation analysis, p < 0.05, n = 10; indicated by cooler colors in DA maps DURING-PRE), and the differences were maintained after stimulation (POST1-PRE and POST2-PRE, Fig. 6A). In this condition, the significant decrease in the spectral power covered a bandwidth between 70-100 Hz (0 to 40 ms) during tDCS application, during 20 minutes after tDCS (POST1-PRE in Fig. 6A) and extended to lower frequencies (40-100 Hz; 0 to 40 ms and 70 to 100 ms) for the last 20 min (POST2-PRE in Fig. 6A). In contrast, during 20 minutes of anodal tDCS intervention the spectral power was significantly higher (permutation analysis, p < 0.05, n = 10, indicated by warm colors in DA maps) with respect to the control period (DURING-PRE in Fig. 6B) covering a bandwidth between 20-50 Hz (0 to 20 ms). Unlike the results of the SEP analysis, where no differences were obtained after anodal tDCS (Fig. 4A,C), we found an increase in the spectral power corresponding to the time frame after whisker stimulation (> 0 ms) at two bandwidths between 30-50 Hz (10 to 50 ms) and 60-100 Hz (0 to 20 ms) throughout the 20 minutes after anodal tDCS (POST1-PRE in Fig. 6B) with only a few significant changes in the following 20 min (POST2-PRE in Fig. 6B). With respect to the time frame previous to the whisker stimulation (< 0 ms) the spectral power decreased in the 30-60 Hz bandwidth (−40 to −20 ms) during anodal tDCS intervention and throughout the 20 min after stimulation, and increased in the 50-80 Hz bandwidth (−20 to 0 ms) in the whole period following tDCS removal (POST1-PRE and POST2-PRE in Fig. 6B). There were no significant changes in the sham group (n = 10) except for small-scattered differences (Fig. 6C).

**Figure 6.**
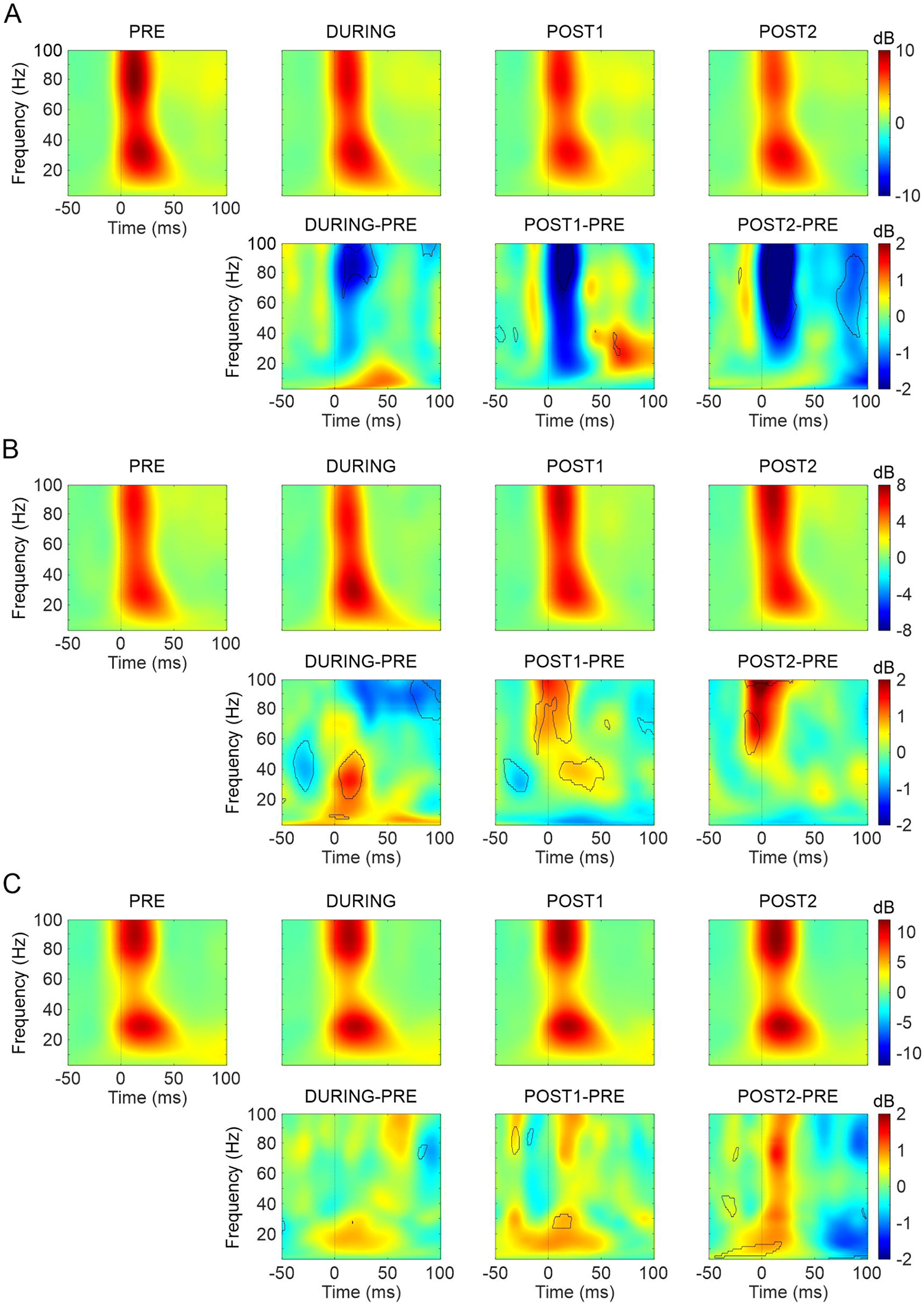
ERSP analysis of the induced activity. *A-C*) (Upper row) ERSP obtained for the 20 min before stimulation (PRE, first column), during tDCS (DURING, second column), the first 20 minutes after tDCS cessation (POST1, third column) and the next 20 min (POST2, fourth column) for cathodal (*A*), anodal (*B*) or sham (*C*) tDCS. (Lower row) Differences between the ERSP obtained in the 20 min before stimulation (PRE) and DURING (first column), POST1 (second column) or POST2 (third column) for cathodal (*A*), anodal (*B*) or sham (*C*) tDCS. Black outline represents statistical differences for that frequency and time range (p < 0.05, permutations).

### Cathodal tDCS induces GAD 65-67 but not vGLUT1 changes in S1 cortex

To elucidate potential molecular expression changes underlying the observed asymmetric long-term effects of anodal vs. cathodal tDCS, we used antibodies against vGLUT1 and GAD 65-67 to assess possible modifications of the excitation/inhibition balance in the transcranially stimulated S1. A group of animals prepared for tDCS application during whisker stimulation (no electrophysiological recordings were carried out in this experiment) was randomly assigned to anodal (n = 5), cathodal (n = 4) or sham (n = 4) condition. Mice from each group were transcardially perfused just after 20 min of anodal, cathodal or sham tDCS and the brains were processed for immunohistological analysis. Representative confocal images from the non-stimulated left hemisphere and the transcranially stimulated right hemisphere are presented in figure 7 for cathodal (at top), anodal (at middle) and sham (at bottom) groups for GAD 65-67 (Fig. 7A) and vGLUT1 (Fig. 7B). The number of GAD 65-67 and vGLUT1 positive clusters of puncta in the stimulated and non-stimulated S1 were analyzed in the cathodal, anodal and sham groups. We obtained a significant general main effect on the GAD 65-67 positive clusters for the interaction BRAIN HEMISPHERE x tDCS POLARITY (two-way mixed ANOVA, F_2,10_ = 5.163, p = 0.029, Fig. 7A) with a significant difference between the stimulated vs. non-stimulated hemisphere in the cathodal tDCS condition (n = 4 animals, Bonferroni, p = 0.005, Fig. 7A) indicating a higher number of GAD 65-67 positive clusters in the stimulated S1 hemisphere than in the non-stimulated S1. There was no significant difference in vGLUT1 between the stimulated and non-stimulated hemisphere in any of the tested stimulation conditions (two-way mixed ANOVA, F_2,10_ = 0.12, p = 0.888, Fig. 7B). No significant differences were found for anodal or sham condition. This result suggests that the decrease of SEPs amplitude and the associated power spectrum decrease observed after cathodal tDCS could be mediated by a GABA level imbalance between the stimulated and nonstimulated hemisphere caused by a GABA reduction in the non-stimulated side. A similar decrease in GABA only observed in non-stimulated primary motor cortex after cathodal tDCS has been previously reported in humans (Bachtiar et al., 2018). Finally, to exclude possible spreading effects of cathodal tDCS in the mouse cortex we also tested GAD 65-67 and vGLUT1 positive clusters in the adjacent primary motor cortex (M1). No significant effects were found in any GAD 65-67 and vGLUT1 positive clusters (two-way mixed ANOVA, F_2,10_ = 0.06, p = 0.942, Fig. 7C) suggesting a focalized histological long-term effect of cathodal tDCS on the stimulated region.

**Figure 7.**
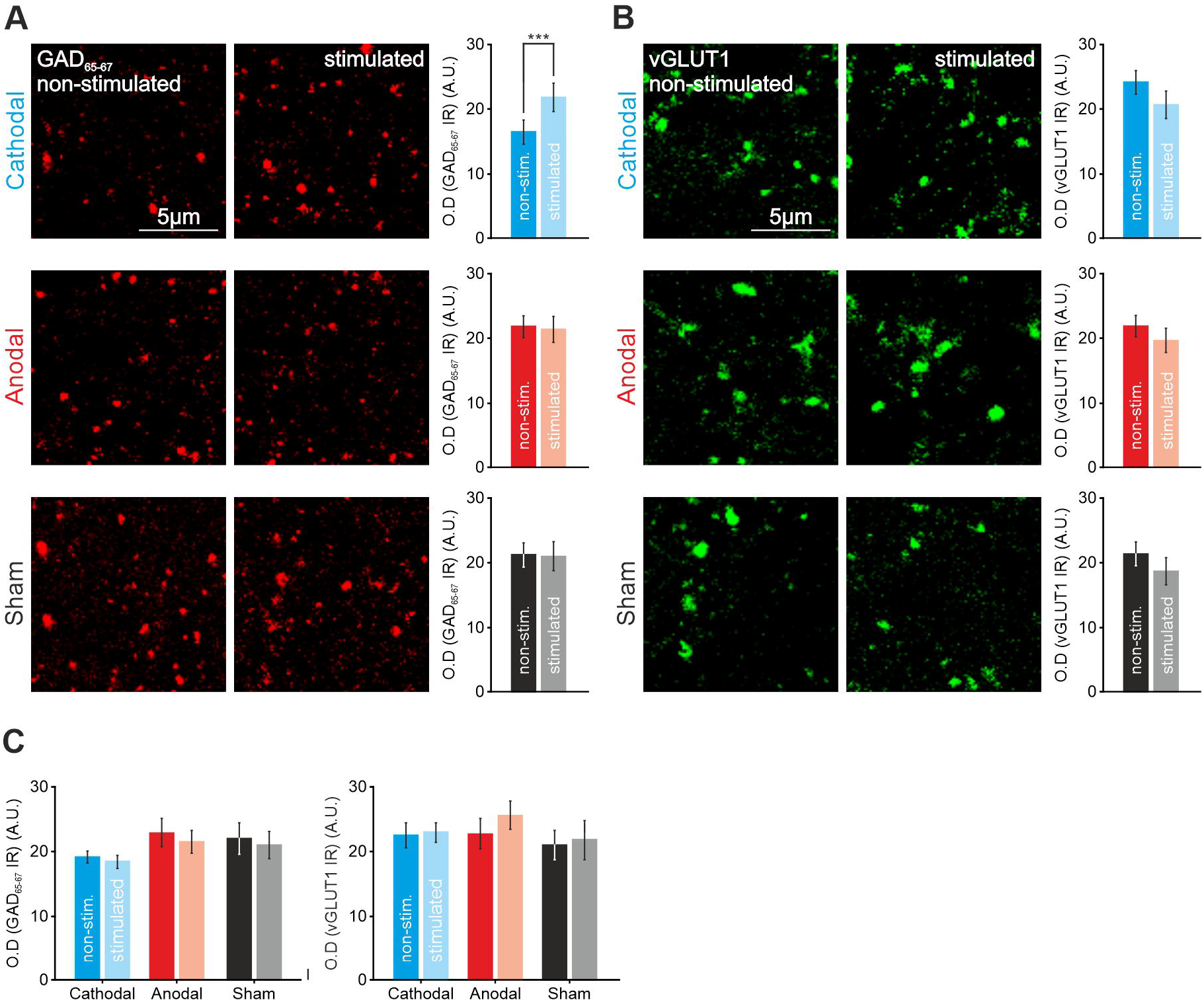
Immunohistochemical changes after 20 minutes of S1-tDCS. *A,B*) Confocal photomicrographs (images), quantification and statistics (bar charts) of GAD 65-67 (*A*) or vGLUT1 immunoreactivity (*B*) in S1 after 20 min of cathodal tDCS (upper row, n = 4 mice, two-way mixed ANOVA, F_2,10_= 5.163, ***p < 0.001), after 20 min of anodal tDCS (middle row, n = 5 mice) and after 20 min of sham condition (lower row, n = 4 mice). *C*) Same analysis but for adjacent motor cortex. Error bars represent SEM. GAD 65-67: Glutamic acid decarboxylase isoforms 65 and 67; vGLUT1: vesicular glutamate transporter 1; OD: optical density; IR: immunoreactivity; A.U.: arbitrary units.

## Discussion

The present electrophysiological and histological results point to asymmetric differences between immediate and long-term effects observed during and after anodal or cathodal S1-tDCS. These results could be of crucial importance for human tDCS protocols suggesting that tDCS polarities could differently impact on cortical excitation/inhibition balance during or after its application.

The first aim of this study was to directly quantify the strength of the induced electric field at different cortical depths. As expected, we observed bigger electric field values in the first millimeter of the cortex following a logarithmic decay with increasing distance. In line with previous studies performed in humans and non-human primates electric fields generated during tES behave in a linear ohmic manner (Opitz et al., 2016). However, in the present study the electric field (23.1 – 90.2 V/m) imposed on the cortical layer of the mice, resulted to be considerably higher than those typically used in humans (1 V/m) (Opitz et al., 2016; Chhatbar et al., 2018). Interestingly, previous investigations showed that electric field intensities used in humans generally fail to generate neuronal modulation when applied to animal models *in vivo* (Ozen et al., 2010; Vöröslakos et al., 2018). A possible explanation for this divergence could be related to differences in axonal lengths (Chakraborty et al., 2018) or larger neuronal densities found in primates (Herculano-Houzel, 2009). In addition, the size of the electrodes used in human experiments, covering large cortical regions, could be optimizing neuronal recruitment by increasing network emergent effects (Fröhlich eta McCormick, 2010; Reato et al., 2010).

On the other hand, the present study took advantage of intracortical LFP and SEP recording in alert mice showing that SEPs, easily identifiable by their waveform, constitute a consistent marker for indexing cortical activity over long periods of time. This method helped us to characterize the electrophysiological changes associated with the immediate and long-term effects of tDCS in S1, an approach that has been successfully used in the past for concurrent DC application and SEP recording in anesthetized rats (Bindman et al., 1964) and tDCS application in alert rabbits (Márquez-Ruiz et al., 2012, 2016). The N1 component of S1-SEPs was particularly susceptible to tDCS. N1 is supposed to represent postsynaptic activity from layers IV, V and VI (Castro-Alamancos and Oldford, 2002; De Kock et al., 2007; Sermet et al., 2019), which explains our observation of a polarity inversion when going deeper than layer VVI.

Our second aim was to investigate immediate effects of tDCS over the S1 mouse cortex. Our results indicate that tDCS applied for several seconds over S1 is sufficient to modulate cortical excitability in agreement with previous results reported from the human motor cortex (Nitsche and Paulus, 2000). We observed an increase of SEP amplitude during anodal stimulation and a decrease during cathodal stimulation. Furthermore, tDCS modulated cortical excitability in an intensity-dependent manner, i.e. greater intensities induced greater changes. Similar results were previously obtained in S1 cortex from rabbits (Márquez-Ruiz et al., 2012, 2016) and in motor (Cambiaghi et al., 2010) and visual (Cambiaghi et al., 2011; Monai et al., 2016) cortices in mice. Overall, these results point toward a similar immediate effect of tDCS across different cortices, at least for the simplified cortical geometry of mice and rabbits where the axo-dendritic orientation of pyramidal cells with respect to the exogenous electric field is homogeneous (Bikson et al., 2004; Radman et al., 2009; Kabakov et al., 2012).

Our third aim was to explore whether tDCS induces long-term changes in SEP amplitude. Unlike immediate effects, a long-lasting modulation of SEP amplitude was only observed after cathodal tDCS, with no changes after anodal tDCS. Similar asymmetric results have been previously reported in alert rabbits where cathodal but not anodal S1-tDCS was able to induce long-term effects measured by a reduction of SEP amplitude after the stimulation (Márquez-Ruiz et al., 2012). Human studies have also reported diverging effects of tDCS. First of all, a few studies have reported a similar absence of effects after anodal tDCS intervention in humans (Rogalewski et al., 2004; Dieckhöfer et al., 2006). Whereas in other studies performed in humans anodal S1-tDCS increased the amplitude of somatosensory evoked magnetic fields (Sugawara et al., 2015) and improved performance in a complex somatosensory task after current application (Ragert et al., 2008). In conformity with our results, reported effects after stimulation in humans indicate that cathodal S1-tDCS besides decreasing tactile perception (Rogalewski et al., 2004), reduced SEP amplitude (Dieckhöfer et al., 2006; Vaseghi et al., 2015) in correlation with increasing sensory and pain thresholds (Vaseghi et al., 2015). The asymmetry of long-term effects observed in the present study is an important issue since long-lasting excitability changes are crucial for clinical treatments (Stagg et al., 2018). Moreover, the lack of long-term effects after anodal stimulation suggests that anodal tDCS may be most effective when applied online, during a given task, rather than before or after it (Ammann et al., 2016; Stagg et al., 2018).

An additional approach of the present study to further explore long-lasting effects of tDCS has been the power spectrum analysis. In this regard, analysis of the power spectrum not in phase with the sensory events (FFT) showed that only cathodal (but not anodal) tDCS affected the amplitude of oscillations (ranging from 20 to 80 Hz) throughout the 20 min after transcranial stimulation. Moreover, sensory event-related spectral dynamics analysis showed that cathodal tDCS was able to decrease the spectral power in a wide range of frequencies (60 to 100 Hz) during the intervention as well as for up to 20 minutes after the stimulation, and in the range between 50-100 Hz for up to 40 minutes after tDCS. On the other hand, anodal tDCS increased the range between 20-50 Hz during stimulation, and 30-50 Hz and 60-100 Hz for up to 20 minutes after the stimulation. Thus, the application of tDCS on S1 seems to modulate gamma activity both during and after transcranial intervention. Accordingly, tDCS intervention in humans has shown to modulate brain oscillations at different frequencies and cortical regions. Specifically, cathodal tDCS caused a significant decrease of spontaneous and induced gamma in the occipital cortex (Antal et al., 2004; Wiesman et al., 2018). On the other hand, anodal tDCS increased spontaneous theta and alpha frequency powers in prefrontal and occipital cortices (McDermott et al., 2019; Wiesman et al., 2018), and induced gamma, beta and gamma in occipital cortices after tDCS (Antal et al., 2004). Interestingly, gamma oscillations have been related to visual attention (Gray et al., 1990; Engel and Singer, 2001), codification, retention and retrieval of information independently of sensory modality (Tallon-Baudry and Bertrand, 1999; Herrmann et al., 2004; Kahana, 2006) together with sensory perception (Siegle et al., 2014). Overall, our finding suggests that tDCS may provide an effective method to modulate a variety of cognitive functions (Berryhill and Martin, 2018).

Finally, to examine potential long-term changes in glutamate and GABA expression associated to anodal and cathodal tDCS we used antibodies against vGLUT1 and GAD 65-67. We observed a significant increase in GAD 65-67 immunoreactivity in the stimulated region respect to the non-stimulated side after cathodal stimulation but no changes for anodal stimulation. This result is in line with our electrophysiological measures, suggesting an overall decrease in the excitability of the stimulated cortex after cathodal tDCS, but no long-lasting effects after anodal tDCS. In humans, polarity-dependent effects on GABA and glutamatergic levels after M1-tDCS have been reported (Stagg et al., 2009), indicating a relation between long-lasting tDCS effects and the cortical excitation/inhibition balance (Krause et al., 2013). Some studies have shown a decrease in GABA after anodal M1-tDCS in the stimulated site (Stagg et al., 2009; Bachtiar et al., 2018; Patel et al., 2019) and in the non-stimulated M1 (Bachtiar et al., 2018). Interestingly enough, Bachtiar and colleagues (2018) describe a decrease in GABA only in the non-stimulated M1 after cathodal stimulation (Bachtiar et al., 2018), similar to what we observed after cathodal tDCS in our experiments.

According to our results, optimal selection of tDCS parameters should be based on extensive knowledge of the brain mechanisms underlying the immediate and longterm impact of exogenous electric fields on the neural network of interest (Jackson et al., 2016; Cirillo et al., 2017; Liu et al., 2018). These heterogeneous mechanisms could explain the reported effects of tDCS in human subjects with different protocol parameters such as the position of stimulating electrodes over the scalp, the polarity, duration and density of the current (Bikson et al., 2019a). Nevertheless, the complication of recording the actual electric field generated inside the human brain, together with the common use of indirect measurements of cortical excitability (Stagg et al., 2018), makes it difficult to examine tDCS-associated mechanisms in human studies.

In summary, the present electrophysiological and immunohistological study clearly shows differences between immediate and long-term tDCS effects on S1, besides a distinct functional asymmetry in anodal and cathodal associated long-term effects. The complexity of the reported effects highlights the importance of defining both the immediate and long-term impact of tDCS on neural processing, to help improve stimulation protocols for treating neurological disease in the clinic.

## Acknowledgements

This work was supported by grants from the Spanish MINECO-FEDER (BFU2014-53820-P and BFU2017-89615-P) to J.M-R and from the US National Institutes of Health (RF1MH114269) to J.F.M and J.M-R. C.A.S-L was in receipt of an FPU grant from the Spanish Government (FPU13/04858).

## Author contributions

C.A.S-L and J.M-R conceived the original idea and designed the experiments. C.A.S-L, I.C, C.A, J.M.A, M.A.G-C and J.M-R performed the experiments and the data analysis. A.C-G and G.S-GC participated in part of the experiments. C.A.S-L and J.M-R wrote the paper. A.G, J.M.D-G, G.C and J.F.M assisted in the experimental design and in the interpretation of the results. All of the authors contributed to the final edition of the manuscript.

## Additional Information

The authors declare no competing financial interests.

